# Dating ancient humans splits by estimating Poisson rates from mitochondrial DNA parity samples

**DOI:** 10.1101/2023.04.21.537838

**Authors:** Keren Levinstein Hallak, Saharon Rosset

## Abstract

We tackle the problem of estimating species divergence times, given a genome sequence from each species and a large known phylogenetic tree with a known structure (typically from one of the species). The number of transitions at each site from the first sequence to the other is assumed to be Poisson distributed, and only the parity of the number of transitions is observed. The detailed phylogenetic tree contains information about the transition rates in each site. We use this formulation to develop and analyze multiple estimators of the divergence between the species. To test our methods, we use mtDNA substitution statistics from the well-established Phylotree as a baseline for data simulation such that the substitution rate per site mimics the real-world observed rates. We evaluate our methods using simulated data and compare them to the Bayesian optimizing software BEAST2, showing that our proposed estimators are accurate for a wide range of divergence times and significantly outperform BEAST2. We then apply the proposed estimators on Neanderthal, Denisovan, and Chimpanzee mtDNA genomes to better estimate their TMRCA (Time to Most Recent Common Ancestor) with modern humans and find that their TMRCA is substantially later, compared to values cited recently in the literature.

## 1 Introduction

Dating species divergence has been studied extensively for the last few decades using approaches based on genetics, archaeological findings, and radiocarbon dating [8, 32]. Finding accurate timing is crucial in analyzing morphological and molecular changes in the DNA, in demographic research, and in dating key fossils. One approach for estimating the divergence times is based on the molecular clock hypothesis [38, 37] which states that the rate of evolutionary change of any specified protein is approximately constant over time and different lineages. Subsequently, statistical inference can be applied to a given phylogenetic tree to infer the dating of each node up to calibration.

Our work focuses on this estimation problem and proposes new statistical methods to date the TMRCA of two species given a detailed phylogenetic tree for one of the species with the same transition rates per site. We formulate the problem by modeling the number of transitions (*A ↔ G, C ↔ T*) in each site using a Poisson process with a different rate per site; sites containing transversions are neglected due to their sparsity (indeed, we include sparse transversions in the simulations and show that our methods are robust to their occurrences). The phylogenetic tree is used for estimating the transition rates per site. Hence, our problem reduces to two binary sequences where the parity of the number of transitions of each site is the relevant statistic from which we can infer the time difference between them.

We can roughly divide the approaches to solve this problem into two. The **frequentist** approach seeks to maximize the likelihood of the observed data. Most notable is the PAML [36] package of programs for phylogenetic analyses of DNA and the MEGA software [18]. Alternatively, the **Bayesian** approach considers a prior of all the problem’s parameters and maximizes the posterior distribution of the observations. Leading representatives of the Bayesian approach are BEAST2 [5] and MrBayes [27], which are publicly available programs for Bayesian inference and model choice across a wide range of phylogenetic and evolutionary models.

In this work, we developed several distinct estimators from frequentist and Bayesian approaches to find the divergence time directly. The proposed estimators differ in their assumptions on the generated data, the approximations they make, and their numerical stability. We explain each estimator in detail and discuss its properties.

A critical difference between our proposed solutions and existing methods is that we seek to estimate only one specific problem parameter. At the same time, software packages such as BEAST2 and PAML optimize over a broad set of unknown parameters averaging the error on all of them (the tree structure, the timing of every node, the per-site substitution rates, etc.). Subsequently, the resources they require for finding a locally optimal instantiation of the tree and dating all its nodes can be very high in terms of memory and computational complexity. Consequently, the amount of sequences they can consider simultaneously is highly limited. Thus, unlike previous solutions, we utilize transition statistics from all available sequences, in the form of a previously built phylogenetic tree.

We develop a novel approach to simulate realistic data to test our proposed solutions. To do so, we employ Phylotree [33] – a complete, highly detailed, constantly updated reconstruction of the human mitochondrial DNA phylogenetic tree. We sample transitions of similar statistics to Phylotree and use it to simulate a sequence at a predefined trajectory from Phylotree’s root.

We then empirically test the different estimators on simulated data and compare our results to the BEAST2 software. Our proposed estimators are calculated substantially faster while utilizing the transitions statistics from all available sequences (Phylotree considers 24,275 sequences), unlike BEAST2 which can consider only dozens of sequences due to its complexity. Comparing with the ground truth, we show that BEAST2 overestimates the divergence time for low TMRCA values (e.g. human-Neanderthals and human-Denisovan), but performs an underestimation for larger divergence times (e.g. human-Chimpanzee), while our estimates provide more accurate results. Finally, we use our estimators to date the TMRCA (given in kya – kilo-years ago) of modern humans with Neanderthals, Denisovan and Chimpanzee based on their mtDNA. Surprisingly, the divergence times we find (human-Neanderthals ∼408 kya, human-Denisovans ∼824 kya, human-Chimpanzee ∼5,009 kya) – are considerably later than those accepted today.

## 2 Materials and Methods

### 2.1 Estimation methods

First, we describe an idealized reduced mathematical formulation for estimating divergence times and our proposed solutions. In Section 2.2, we describe the reduction process in greater detail.

Consider the following scenario: we have a set of *n* Poisson rates, denoted as 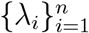 where *n* ∈ ℕ. Let 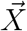 be a vector of length *n* such that each element *X*_*i*_ is independently distributed as Pois(*λ*_*i*_). Similarly, let 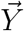 be a vector of length *n* such that each element *Y*_*i*_ is independently distributed as Pois(*λ*_*i*_ *·p*) for a fixed unknown *p*. We denote 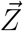 as the coordinate-wise parity of 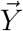, meaning that *Z*_*i*_ = 1 if *Y*_*i*_ is odd and *Z*_*i*_ = 0 otherwise. **Our goal is to estimate** *p* **given** 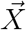 **and** 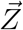.

#### Remark 1

Note that the number of *unknown* Poisson rate parameters *n* in the problem 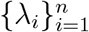 grows with the number of observations 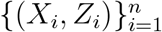. However, our focus is solely on estimating *p*, so additional observations do provide more information.

#### Remark 2

The larger the value of *p ·λ*_*i*_, the less information on *p* is provided in *Z*_*i*_ as it approaches a Bernoulli distribution with a probability of 0.5. On the other hand, the smaller *λ*_*i*_ is, the harder it will be to infer *λ*_*i*_ from *X*_*i*_. As a result, the problem of estimating *p* should be easier in settings where *λ*_*i*_ is high and *p* is low.

#### 2.1.1 Preliminaries

First, we derive the distribution of *Z*_*i*_; All proofs are provided in the Supplementary material (Section 1).

##### Lemma 1.

*Let Y* ∼ *Pois*(Λ) *and Z be the parity of Y*. *Then* 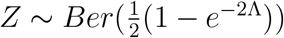.

We use this result to calculate the likelihood and log-likelihood of *p* and 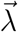 given 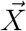 and 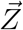. The likelihood is given by:

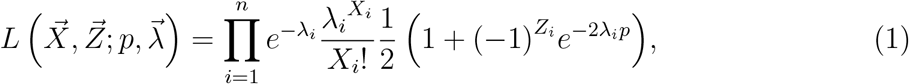

and the log-likelihood is:

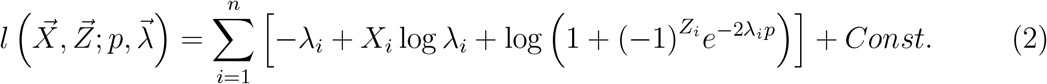

This result follows immediately from the independence of each coordinate.

#### 2.1.2 Cramer-Rao bound

We begin our analysis by computing the Cramer-Rao bound (CRB; [7, 25]). In Section 3.1, we compare the CRB to the error of the estimators.

##### Theorem 1.

*Denote the Fisher information matrix for the estimation problem above by I* ∈ *ℝ* ^(*n*+1,*n*+1)^, *where the first n indexes correspond to* 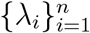 *and the last index* (*n* + 1) *corresponds to p. For clarity denote I*_*p,p*_ ≐ *I*_*n*+1,*n*+1,_*I*_*i,p*_ ≐ *I*_*i,n*+1_, *I*_*p,i*_ ≐ *I*_*n*+1,*i*_. *Then:*

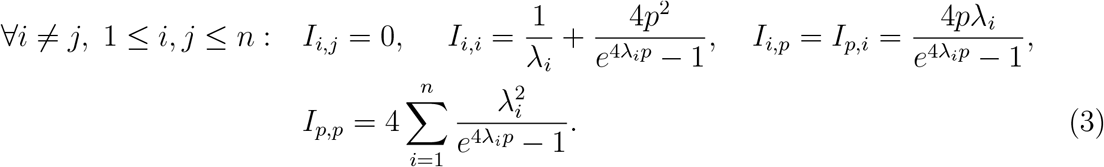

*Consequently, an unbiased estimator* 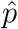 *holds:*

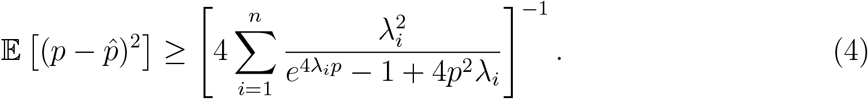

*If* ∀*i* = 1..*n* : *λ*_*i*_ = *λ, we can further simplify the expression:*

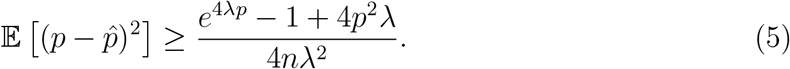

The CRB, despite its known looseness in many problems, provides insights into the sensitivity of the error to each parameter. This expression supports our previous observation that the error of an unbiased estimator increases exponentially with min_*i*_{*λ*_*i*_ *·p*}. However, for constant *λ*_*i*_ *·p*, the error improves for higher values of *λ*_*i*_. We now proceed to describe and analyze several estimators for *p*.

#### 2.1.3 Method 1 - Maximum Likelihood Estimator

##### Proposition 1.

*Following equation 1, the maximum likelihood estimators* 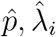 *hold:*

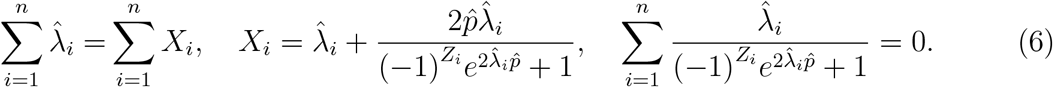

**Proposition 1** provides *n* separable equations for maximum likelihood estimation (MLE). Our first estimator sweeps over values of 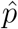 (grid searching in a relevant area) and then for each *i* = 1..*n* finds the optimal 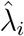 numerically. The solution is then selected by choosing the pair 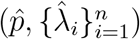 that maximizes the log-likelihood calculated using equation 2.

The obtained MLE equations are solvable, yet, finding the MLE still requires solving *n* numerical equations, which might be time-consuming. More importantly, MLE estimation is statistically problematic when the number of parameters is of the same order as the number of observations [6]. Subsequently, we propose alternative methods that may yield better practical results.

#### 2.1.4 Method 2 - *λ*_*i*_-conditional estimation

We propose a simple estimate of 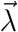 based solely on *X*_*i*_, followed by an estimate of *p* as if 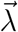 is known, considering only 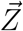. This method is expected to perform well when *λ*_*i*_ values are large, as in these cases, *X*_*i*_ conveys more information about *λ*_*i*_ than *Z*_*i*_. This approach enables us to avoid estimating both 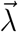 and *p* simultaneously, leading to a simpler numerical solution.

When *p* ≤ 1, we can mimic *Y*_*i*_’s distribution as a sub-sample from *X*_*i*_, i.e. we assume that *Y*_*i*_|*X*_*i*_ ∼ *Bin*(*n* = *X*_*i*_, *p*). Then, we find the maximum likelihood estimate of *p*:

##### Proposition 2.

*If Y*_*i*_|*X*_*i*_ ∼ *Bin*(*X*_*i*_, *p*), *then:*

1. *Y*_*i*_ ∼ *Pois*(*λ*_*i*_ *·p*), *which justifies this approach*.
2. 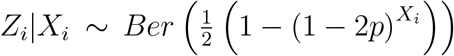, *so we can compute the likelihood of p without considering λ*_*i*_.
3. *The maximum likelihood estimate of p given* 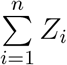 *holds:*

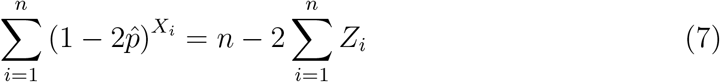

**Remark:** We use the maximum likelihood estimation of *p* given 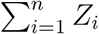 by applying Le-Cam’s theorem [20]. This eliminates the need for a heuristic solution of the pathological case *X*_*i*_ = 0, *Z*_*i*_ = 1.

#### 2.1.5 Method 3 - Gamma distributed Poisson rates

The Bayesian statistics approach incorporates prior assumptions about the parameters. A common prior for the rate parameters 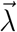 is the Gamma distribution, which is used in popular Bayesian divergence time estimation programs such as MCMCtree [36], BEAST2 [5], and MrBayes [27]. Specifically, we have *λ*_*i*_ ∼ Γ(*α, β*), and for *p*, we use a uniform prior over the positive real line.

##### Proposition 3.

*Let λ*_*i*_ ∼ Γ(*α, β*), *then the maximum a posteriori estimator of p holds:*

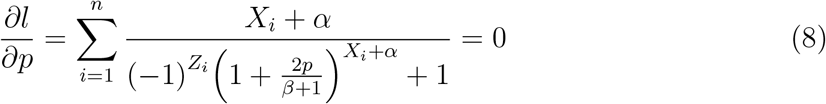

Subsequently, given estimated values for *α* and *β*, we can be find an estimator for *p* numerically to hold Equation 8. Unfortunately, the derivative with respect to *α* does not have a closed-form expression, nor is it possible to waive the dependence on 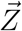, *p*. Hence, we suggest using Negative-Binomial regression [12] to estimate *α* and *β* given 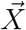.

### 2.2 Estimating ancient divergence times using a large modern phylogeny

In this section, we apply the methods described in Section 2.1 to estimate the non-calibrated divergence times between humans and their closest relatives by comparing mitochondrial DNA (mtDNA) sequences. Our approach assumes the following assumptions:

1. Molecular clock assumption - the rate of accumulation of transitions (base changes) over time and across different lineages is constant, as first proposed by Zuckerkandl and Pauling [38] and widely used since.
2. Poisson distribution - The number of transitions along the human and human’s closest relatives mtDNA lineages follows a Poisson distribution with site-dependent rate parameter *λ*_*i*_ per time unit.
3. No transversions - We only consider sites with no transversions and assume a constant transition rate per site (*λ*_*i,A*→*G*_ = *λ*_*i,G*→*A*_, or *λ*_*i,T* →*C*_ = *λ*_*i,C*→*T*_).
4. Independence of sites - The number of transitions at each site is independent of those at other sites.
5. Phylogenetic tree - The phylogenetic tree presented in the Phylotree database includes all transitions and transversions that occurred along the described lineages.

As the Phylotree database is based on tens of thousands of sequences, the branches in the tree correspond to relatively short time intervals, making multiple mutations per site unlikely in each branch [29]. However, when considering the mtDNA sequence of other species, the branches in the tree correspond to much longer time intervals, meaning that many underlying transitions are unobserved. For instance, when comparing two human sequences that differ in a specific site, Phylotree can determine whether the trajectory between the sequences was *A* → *G, A* → *G* → *A* → *G*, or *A* → *T* → *G*. However, when comparing sequences of ancient species, an elaborate phylogenetic tree like Phylotree is not available, making it impossible to discriminate between these different trajectories.

We use the following notation:

1. Let 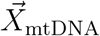 denote the number of transitions observed at each site along the human mtDNA phylogenetic tree as described by Phylotree. Each coordinate corresponds to a different site out of the 16,569 sites. The number of transitions at site *i, X*_mtDNA,i_, follows a Poisson distribution with parameter *λ*_*i*_.
2. Let 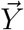 denote the number of transitions between two examined sequences (e.g. modern human and Neanderthal). We normalize the length of the tree edges so that the sum of all Phylotree’s edges is one. The estimated parameter *p* relates to the edge distance between the two examined sequences. Subsequently, *Y*_*i*_ follows a Poisson distribution with parameter *λ*_*i*_ *·p*.
3. Let 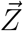 denote the parity of 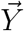.

Using 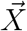 and 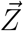, we can estimate *p* using the methods in Section 2.1. The TMRCA is given by: 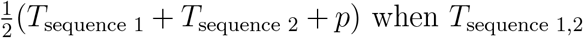 are the estimated times of the examined sequences measured in (uncalibrated) units of phylotree’s total tree length.

### 2.3 Calibration

Our methods output *p*, which is the ratio of two values:

1. The sum of the edges between the two examined sequences and their most recent common ancestor (MRCA).
2. The total sum of Phylotree’s edges.

Similarly to BEAST2, to calibrate *p* to years, we use the per-site per-year substitution rate for the coding region given in [10] *μ* = 1.57 × 10E-8. We then calculate the total sum of Phylotree’s edges in years by dividing the average number of substitutions in the coding region per site (1.4) by *μ*.

### 2.4 Data Availability Statement

The code used in this work is available at: https://github.com/Kerenlh/DivergenceTimes. A full description of all simulations is available in the Materials and Methods section, pages 8-10, and in the Supplementary Material, pages 26-28.

## 3 Results

### 3.1 Comparative Study on Raw Simulations

To compare the performance of the three estimation methods described in 2.1, we conducted experiments using simulated data. The Poisson rates *λ* were generated to reflect the substitution rates observed in mtDNA data using either a Categorical or a Gamma distribution. The parameters for the Gamma distribution (*α* = 0.23, *β* = 0.164) were estimated directly from the data, while the parameters for the Categorical distribution were chosen such that both distributions have the same mean and variance. One of the Categorical values (*E* = 0.1) corresponds to the rate of low activity sites in the mtDNA data. The other value (*a* = 11.87) and the probabilities (0.11, 0.89) were chosen accordingly. The results of the comparison are shown in Figure 1 with the Cramer-Rao bound for reference. To provide a qualitative comparison, we performed a one-sided paired Wilcoxon signed rank test on every pair of models, correcting for multiple comparisons using the Bonferroni correction. Our results show that Method 2 has the lowest squared error while Method 1 has the highest squared error, for both distributions. It is noteworthy that although Method 3 assumes a Gamma distribution, it still performs well even when there is a model mismatch.

**Figure 1:**
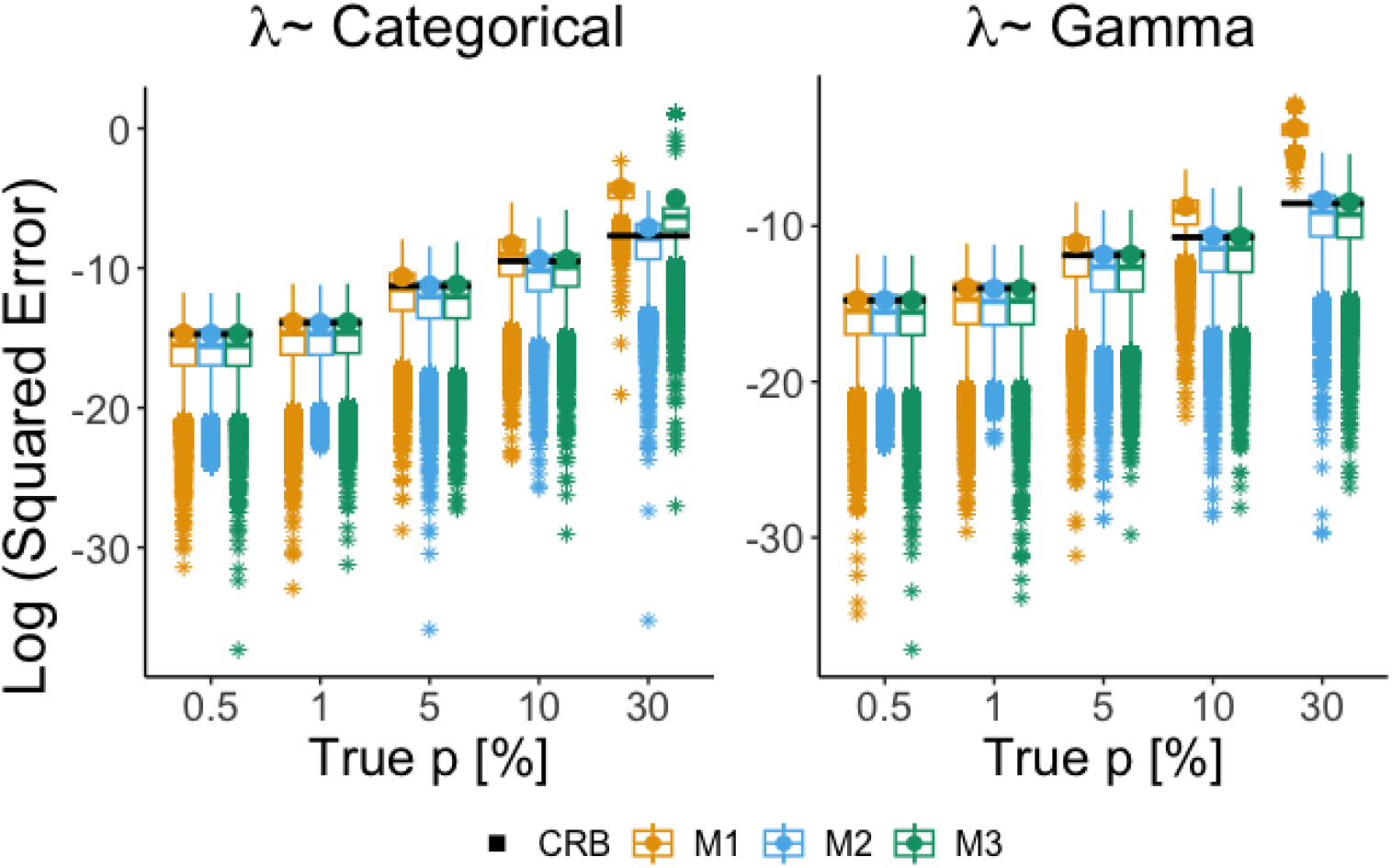
Estimation errors for different *λ* distributions. Box-plot of the log squared estimation errors of the three proposed methods for selected values of *p*, expressed as percentage of the total length of Phylotree’s edges (outliers are marked with *). The simulations were run 10, 000 times for each value of *p*. The CRB is shown in black for reference and the circles represent the log of the mean values which are comparable to the CRB. The experiments were conducted for two different distributions of *λ*: (Left) Categorical distribution with two values: *E* = 0.1 with probability *η* = 0.11 and *a* = 11.87 with probability 1 − *η*. (Right) Gamma distribution with parameters *α* and *β*.

### 3.2 Phylogenetic Tree Simulations

We validated our methods by testing their performance in a more realistic scenario of simulating a phylogenetic tree. Our methods take as input the observed transitions along Phylotree 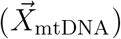 and a binary vector 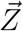 denoting the differences between two sequences, which we aim to estimate the distance between. We compared our methods to the wellknown BEAST2 software [5], which, similarly to other well-established methods (such as MCMCtree [36], MrBayes [27], etc.) considers sequences along with their phylogenetic tree to produce time estimations. The software BEAST2 performs Bayesian analysis using MCMC to average over the space of possible trees. However, it is limited in its computational capacity, so it cannot handle a large number of sequences like those in Phylotree. For this reason, we used a limited set of diverse sequences, including mtDNA genomes of 53 humans [13], the revised Cambridge Reference Sequence (rCRS) [2], the root of the human phylogenetic mtDNA tree, termed Reconstructed Sapiens Reference Sequence (RSRS) [4], and 10 ancient modern humans [10]. More details about the parameters used by BEAST2 are available in the Supplementary material, Section 2.2. To evaluate our methods, we added a simulated sequence with a predefined distance from the RSRS.

Our aim is to generate a vector 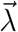 that produces a vector 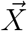 that has a similar distribution to 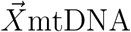. The human mtDNA tree has 16,569 sites, of which 15,629 have no transversions. The MLE of *λ*_*i*_ at each site is the observed number of transitions, *X*_mtDNA,i_. However, simulating 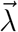 as 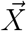 leads to an undercount of transitions because 10,411 sites (67% of the total number of sites considered) had no transitions along the tree and their Poisson rate is taken to be zero. To mitigate this issue, the rates for these sites were chosen to be *E*, the value that minimizes the Kolmogorov-Smirnov statistic [16, 28] (details are provided in the Supplementary material, Section 2.1).

The results are presented in Figure 2. BEAST2 overestimates the true *p* when *p* is smaller than approximately 2%, and underestimates it when *p* is higher. Additionally, BEAST2 has a much longer running time (roughly 3 hours) compared to our methods (less than a second). As shown in Figure 1, Method 1 has a larger error than Methods 2 and 3 for values of *p* within the simulated region, and the gap widens with increasing *p*. Methods 2 and 3 provide the best results for the entire range of p.

**Figure 2:**
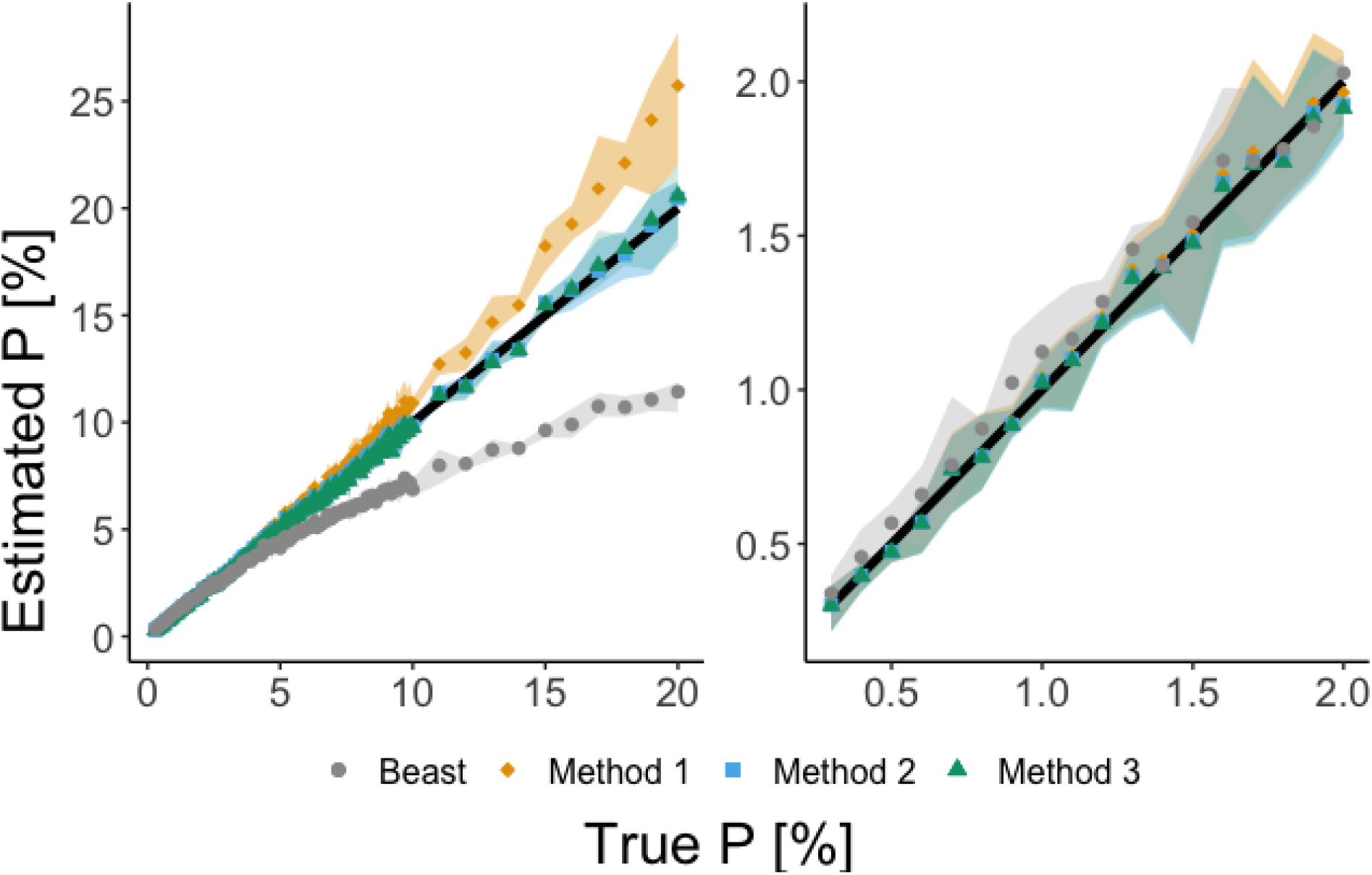
Comparison of estimators applied on a simulated long branch. Comparison of our methods with BEAST2 estimator using simulated data. The right plot shows a zoom-in view of the left plot, focusing on values of *p* between 0 and 10%. Each point in the plot represents the average of 5 runs, while the shaded regions indicate the range of estimations obtained.

### 3.3 Real data results

As the final step of our experiments, we apply our methods on real-world data to determine the TMRCA of the modern human and Neanderthal, Denisovan, and chimpanzee mtDNA genomes. Table 1 displays the uncalibrated distances between modern human and each sequence, compared to the estimates from BEAST2. The presented TMRCA represents an average of the TMRCA obtained from 55 modern human mtDNA sequences of diverse origins [13]. Table 2 presents the TMRCA in kya (kilo-years ago) of the modern human and each sequence.

**Table 1:** Uncalibrated distances between modern humans and selected hominins. Uncalibrated distances expressed as a percentage of the total length of Phylotree’s edges, as determined by our methods compared with BEAST2. The values correspond to *p*, and indicate the estimation’s location in Figure 2. In the parentheses we provide the standard deviation for each estimator, obtained from bootstrapping 100 site samples for every modern human – ancient sequence pair in the dataset. Note that the BEAST2 values presented here were de-calibrated as described in Section 2.3.

| Sample | BEAST2 | Method 1 | Method 2 | Method 3 |
| --- | --- | --- | --- | --- |
| **Altai** | 0.97 ($\pm 0.07$) | 0.8 ($\pm 0.08$) | 0.79 ($\pm 0.08$) | 0.79 ($\pm 0.08$) |
| **Denisova15** | 0.97 ($\pm 0.07$) | 0.81 ($\pm 0.08$) | 0.8 ($\pm 0.08$) | 0.8 ($\pm 0.08$) |
| **HST** | 0.97 ($\pm 0.07$) | 0.78 ($\pm 0.08$) | 0.78 ($\pm 0.08$) | 0.77 ($\pm 0.08$) |
| **Mezmaiskaya1** | 1.01 ($\pm 0.07$) | 0.86 ($\pm 0.09$) | 0.85 ($\pm 0.09$) | 0.85 ($\pm 0.09$) |
| **Chagyrskaya08** | 1.03 ($\pm 0.07$) | 0.84 ($\pm 0.09$) | 0.83 ($\pm 0.09$) | 0.83 ($\pm 0.08$) |
| **ElSidron1253** | 1.05 ($\pm 0.07$) | 0.83 ($\pm 0.08$) | 0.82 ($\pm 0.08$) | 0.82 ($\pm 0.08$) |
| **Vindija33.17** | 1.06 ($\pm 0.07$) | 0.86 ($\pm 0.09$) | 0.85 ($\pm 0.09$) | 0.85 ($\pm 0.09$) |
| **Feldhofer1** | 1.07 ($\pm 0.07$) | 0.85 ($\pm 0.09$) | 0.84 ($\pm 0.08$) | 0.83 ($\pm 0.08$) |
| **GoyetQ56-1** | 1.08 ($\pm 0.07$) | 0.88 ($\pm 0.09$) | 0.88 ($\pm 0.09$) | 0.87 ($\pm 0.09$) |
| **GoyetQ57-2** | 1.08 ($\pm 0.07$) | 0.84 ($\pm 0.09$) | 0.83 ($\pm 0.08$) | 0.83 ($\pm 0.08$) |
| **Les Cottes Z4-1514** | 1.08 ($\pm 0.07$) | 0.91 ($\pm 0.09$) | 0.91 ($\pm 0.09$) | 0.9 ($\pm 0.09$) |
| **Mezmaiskaya2** | 1.07 ($\pm 0.07$) | 0.84 ($\pm 0.09$) | 0.83 ($\pm 0.09$) | 0.83 ($\pm 0.08$) |
| **Vindija33.16** | 1.07 ($\pm 0.07$) | 0.87 ($\pm 0.09$) | 0.86 ($\pm 0.09$) | 0.86 ($\pm 0.09$) |
| **Vindija33.25** | 1.07 ($\pm 0.07$) | 0.85 ($\pm 0.09$) | 0.84 ($\pm 0.09$) | 0.83 ($\pm 0.09$) |
| **GoyetQ305-7** | 1.08 ($\pm 0.07$) | 0.89 ($\pm 0.09$) | 0.89 ($\pm 0.09$) | 0.88 ($\pm 0.09$) |
| **GoyetQ374a-1** | 1.08 ($\pm 0.07$) | 0.89 ($\pm 0.09$) | 0.89 ($\pm 0.09$) | 0.88 ($\pm 0.09$) |
| **Spy 94a** | 1.08 ($\pm 0.07$) | 0.88 ($\pm 0.09$) | 0.88 ($\pm 0.09$) | 0.87 ($\pm 0.09$) |
| **Sima de los Huesos** | 1.7 ($\pm 0.09$) | 1.42 ($\pm 0.12$) | 1.39 ($\pm 0.11$) | 1.39 ($\pm 0.11$) |
| **Denisova2** | 1.88 ($\pm 0.1$) | 1.68 ($\pm 0.13$) | 1.65 ($\pm 0.12$) | 1.64 ($\pm 0.12$) |
| **Denisova8** | 1.92 ($\pm 0.1$) | 1.69 ($\pm 0.13$) | 1.66 ($\pm 0.13$) | 1.65 ($\pm 0.12$) |
| **Denisova4** | 2 ($\pm 0.1$) | 1.83 ($\pm 0.14$) | 1.79 ($\pm 0.13$) | 1.78 ($\pm 0.13$) |
| **Denisova3** | 2.01 ($\pm 0.1$) | 1.82 ($\pm 0.13$) | 1.78 ($\pm 0.13$) | 1.77 ($\pm 0.13$) |
| **Chimpanzee** | 8.31 ($\pm 0.25$) | 12.75 ($\pm 0.68$) | 11.21 ($\pm 0.53$) | 11.21 ($\pm 0.53$) |

**Table 2:** Estimated divergence times between modern human and selected hominins. The table displays the estimated divergence times (in kya) between modern humans and selected hominins, as determined by our methods and compared with BEAST2. The standard deviation, which arises from a combination of the standard deviation of our methods and the sample dating, is given in parentheses. It’s important to note that BEAST2 calculates the TMRCA for all sequences in the same clade as a single estimate, while our methods estimate the TMRCA for each sample individually by taking the average of estimations derived from comparing the sample with every modern human sequence in the dataset.

| Sample | BEAST2 | Method 1 | Method 2 | Method 3 |
| --- | --- | --- | --- | --- |
| Altai | | 426.68 ($\pm 39.79$) | 424.44 ($\pm 39.36$) | 422.02 ($\pm 38.91$) |
| Denisova15 | | 428.8 ($\pm 39.25$) | 426.32 ($\pm 38.78$) | 423.94 ($\pm 38.41$) |
| HST | | 418.96 ($\pm 40.46$) | 416.45 ($\pm 40.02$) | 414.49 ($\pm 39.62$) |
| Mezmaiskaya1 | | 432.86 ($\pm 41.1$) | 430.43 ($\pm 40.57$) | 427.94 ($\pm 40.26$) |
| Chagyrskaya08 | | 416.04 ($\pm 39.86$) | 413.51 ($\pm 39.29$) | 411.02 ($\pm 39$) |
| ElSidron1253 | | 403.29 ($\pm 38.09$) | 400.85 ($\pm 37.58$) | 398.34 ($\pm 37.23$) |
| Vindija33.17 | | 410.76 ($\pm 39.73$) | 408.25 ($\pm 39.18$) | 405.86 ($\pm 38.85$) |
| Feldhofer1 | | 399.36 ($\pm 38.45$) | 396.93 ($\pm 37.91$) | 394.38 ($\pm 37.59$) |
| GoyetQ56-1 | | 416.06 ($\pm 39.76$) | 413.6 ($\pm 39.11$) | 411.1 ($\pm 38.86$) |
| GoyetQ57-2 | | 394.71 ($\pm 38.28$) | 392.29 ($\pm 37.78$) | 389.72 ($\pm 37.43$) |
| Les Cottes Z4-1514 | | 428.9 ($\pm 41.04$) | 426.3 ($\pm 40.42$) | 423.95 ($\pm 40.07$) |
| Mezmaiskaya2 | | 396 ($\pm 38.64$) | 393.59 ($\pm 38.11$) | 391.03 ($\pm 37.74$) |
| Vindija33.16 | | 409.84 ($\pm 39.4$) | 407.47 ($\pm 38.81$) | 404.79 ($\pm 38.53$) |
| Vindija33.25 | | 399.83 ($\pm 39.26$) | 397.4 ($\pm 38.73$) | 394.84 ($\pm 38.39$) |
| GoyetQ305-7 | | 418.12 ($\pm 40.35$) | 415.41 ($\pm 39.69$) | 413.27 ($\pm 39.4$) |
| GoyetQ374a-1 | | 418.12 ($\pm 39.72$) | 415.41 ($\pm 39.06$) | 413.27 ($\pm 38.8$) |
| Spy 94a | | 415.03 ($\pm 40.47$) | 412.57 ($\pm 39.83$) | 410.07 ($\pm 39.55$) |
| Humans-Neandertals | 501.87 ($\pm 31.01$) | 410.91 ($\pm 7.19$) | 408.48 ($\pm 7.09$) | 406.12 ($\pm 7.03$) |
| Sima de los Huesos | | 808.29 ($\pm 61.42$) | 797.88 ($\pm 60.04$) | 795 ($\pm 59.58$) |
| Denisova2 | | 846.56 ($\pm 59.86$) | 831.83 ($\pm 57.75$) | 828.27 ($\pm 57.4$) |
| Denisova8 | | 831.07 ($\pm 61.34$) | 815.67 ($\pm 59.03$) | 812.62 ($\pm 58.78$) |
| Denisova4 | | 857.56 ($\pm 62.64$) | 840.36 ($\pm 60.45$) | 835.84 ($\pm 60$) |
| Denisova3 | | 850.7 ($\pm 60.85$) | 833.74 ($\pm 58.65$) | 829.05 ($\pm 58.25$) |
| Humans-Denisovans-Sima | 934.12 ($\pm 46.54$) | 838.84 ($\pm 27.38$) | 823.9 ($\pm 26.47$) | 820.16 ($\pm 26.3$) |
| Humans-Chimpanzee | 3,711.79 ($\pm 112.13$) | 5,693.51 ($\pm 302.59$) | 5,009.78 ($\pm 235.05$) | 5,005.39 ($\pm 237.13$) |

The estimates from real-world sequences presented in Table 1 are consistent with those obtained for the simulated dataset in Section 3.2. For low values of *p*, our three methods all produce similar estimates while BEAST2’s has a slightly higher estimate. For the human-Chimpanzee uncalibrated distance, which is relatively high, Method 1 provides a higher estimate than that obtained by Methods 2 and 3, while BEAST2 provides a substantially lower estimate. The results in Table 2 show the TMRCA estimates, which are significantly smaller for our methods than those obtained from BEAST2 for human-Neanderthals and human-Denisovans. For example, BEAST2 estimated the human – Sima de los Huesos – Denisovans divergence time as ∼ 934 kya, while our best-performing method (2) estimated it as ∼ 824 kya. This divergence time is estimated as (540-1,410 kya) in [22]. Similarly, BEAST2 estimated the human – Neanderthal divergence time as ∼ 502 kya, while our methods estimated it as ∼ 408 kya. Preceding literature estimates this time closer to ours (∼400 kya [23, 9, 26]) while recent literature provides a much earlier estimate (∼800 kya [11]). Finally, BEAST2 estimates the human-Chimpanzee TMRCA as ∼3,712 kya whereas our estimate is ∼5,001 kya, much closer to the literature value of 5 – 8 million years ago [17, 19, 1, 30].

## 4 Conclusion

We investigated an estimation problem arising in statistical genetics when estimating divergence times between species. The problem’s formulation, estimating Poisson rates from parity samples, leads to multiple estimators with varying assumptions. We calculated the CRB for this estimation problem and compared our methods against commonly used BEAST2 in different empirical settings, including a simple sampling scheme (Section 3.1), a more elaborate generative scheme based on real-world mtDNA data (Section 3.2), and the calculation of the TMRCA of modern humans and other hominins using their mtDNA genomes (Section 3.3).

Our results indicate that our proposed methods are significantly faster and more accurate than BEAST2, especially for earlier divergence times such as the human-Chimpanzee. Our methods utilize the transition statistics from the entire known human mtDNA phylogenetic tree (Phylotree) without the need for reconstructing a tree containing the sequences of interest. Our results show that the human – Neanderthal divergence time is ∼ 408, 000 years ago, considerably later than the values obtained by BEAST2 (∼ 502, 000 years ago) and other values cited in the literature.

## Supplementary Material

### 1 Theoretical Details

#### 1.1 Proof of Lemma 1

Let *Y* ∼ Pois(*λ*) and *Z* be the parity of *Y*. Then 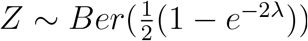.

*Proof*.

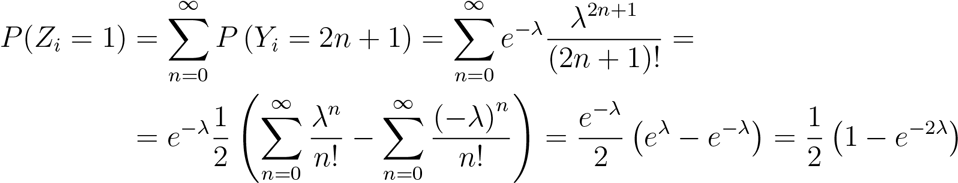

#### 1.2 Proof of Theorem 1

Denote the Fisher information matrix for the estimation problem above by *I* ∈ *ℝ* ^(*n*+1,*n*+1)^, where the first *n* indexes correspond to 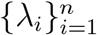 and the last index (*n* + 1) corresponds to *p*. For clarity denote *I*_*p,p*_ ≐ *I*_*n*+1,*n*+1_, *I*_*i,p*_ ≐ *I*_*i,n*+1_, *I*_*p,i*_ ≐ *I*_*n*+1,*i*_. Then:

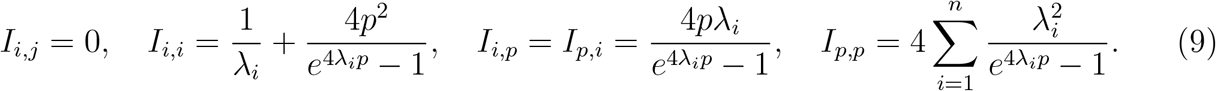

Consequently, an unbiased estimator 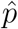 holds:

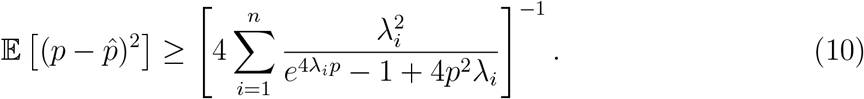

If ∀*i* = 1..*n* : *λ*_*i*_ = *λ*, we can further simplify the expression:

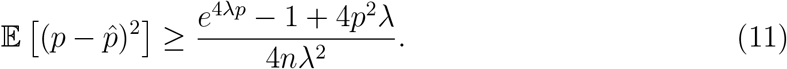

*Proof*. We calculate the second derivative of the log-likelihood. Denote:

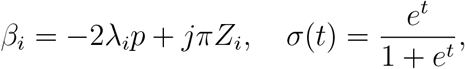

then the first derivatives are given by:

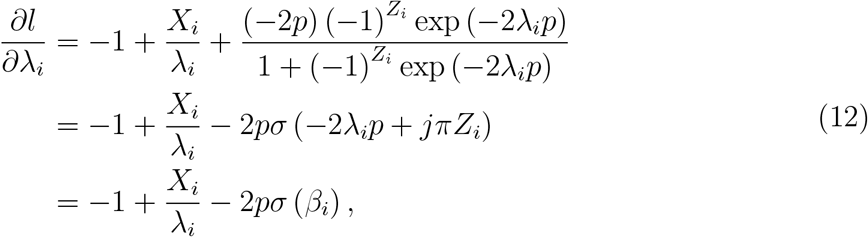

and

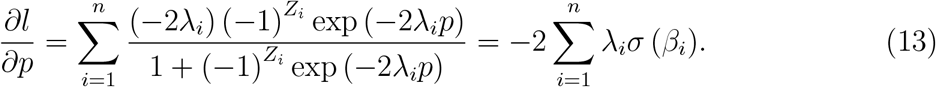

The second derivatives are now given by:

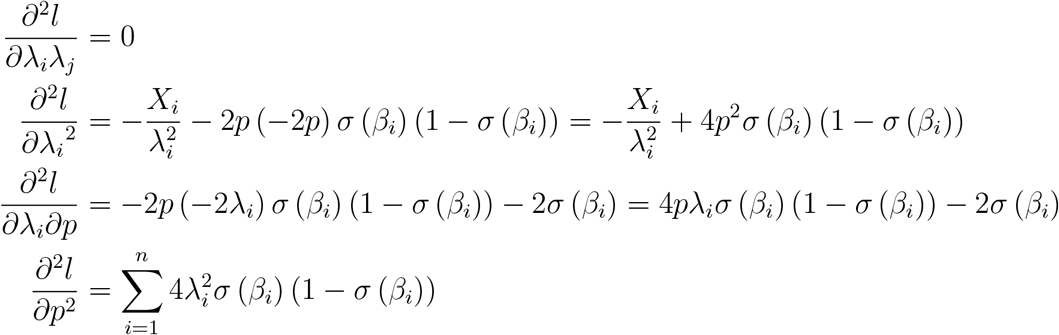

The expectation of these are given by:

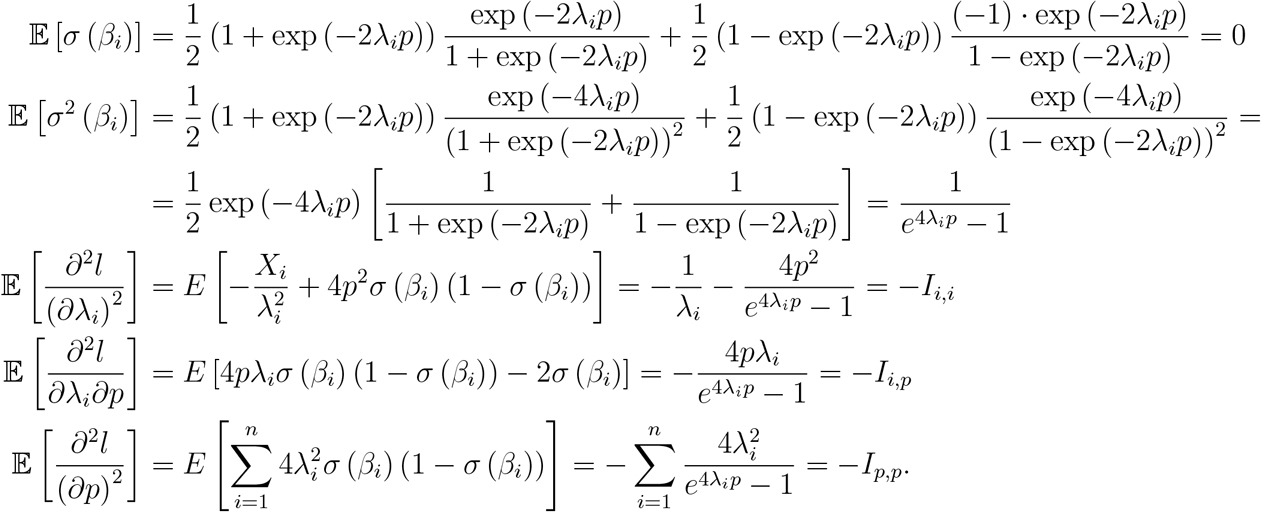

By CRB, for an unbiased estimator:

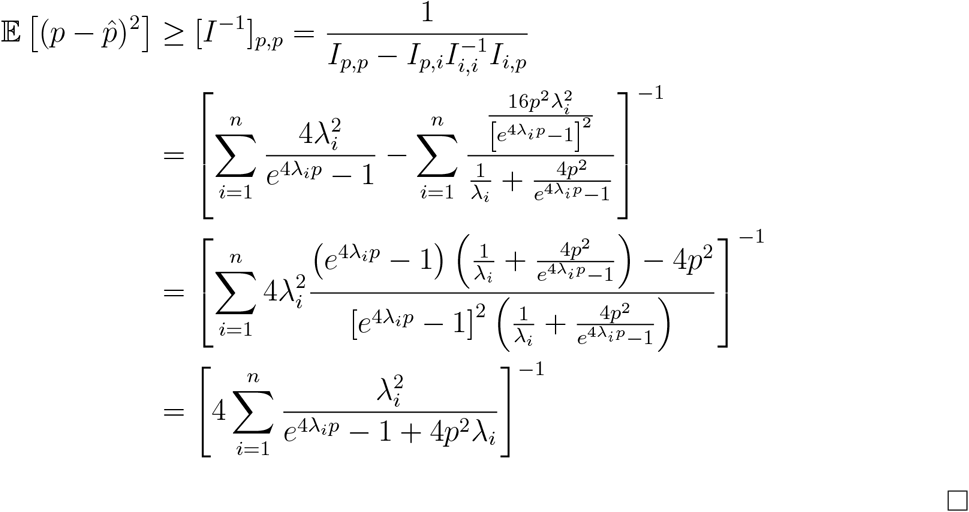

#### 1.3 Proof of Proposition 1

*Proof*. Following Equations 12, 13, we compare the first order derivatives to 0:

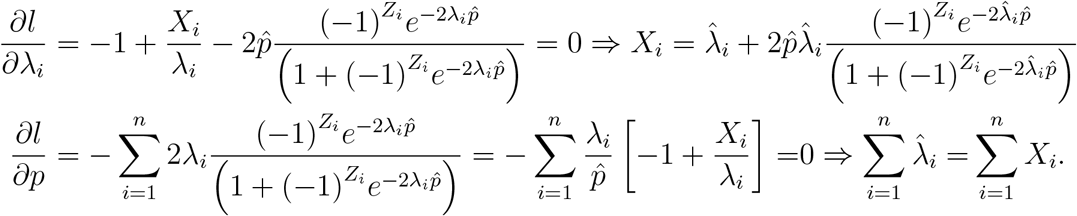

Summing the first equation for every *i* and substituting the second equation results in the last part in Equation 6.

#### 1.4 Proof of Proposition 2

If *Y*_*i*_|*X*_*i*_ ∼ *Bin*(*X*_*i*_, *p*), then:

1. *Y*_*i*_ ∼ Pois(*λ*_*i*_ *·p*), which justifies this approach.
2. 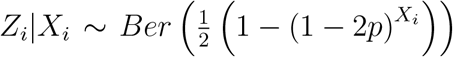, so we can compute the likelihood of *p* without considering *λ*_*i*_.
3. The maximum likelihood estimate of *p* given *Z*_*i*_ holds:

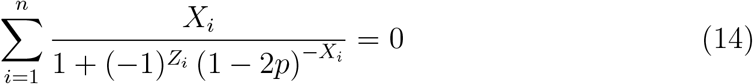

and the maximum likelihood estimate of *p* given 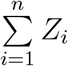 holds:

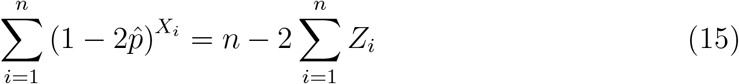

*Proof*. Denote *q* ≡ 1 − *p*. For item 1:

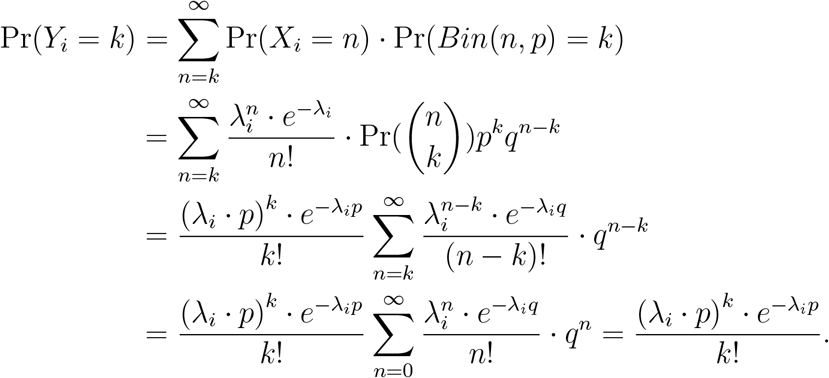

Now moving on to item 2:

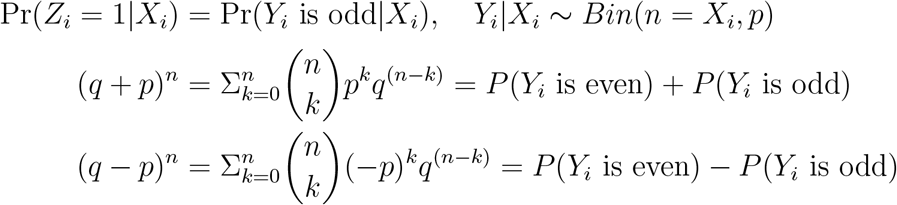

And summing up these two equations leads to:

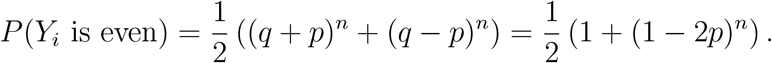

Subsequently, the likelihood of *Z*_*i*_ is given by:

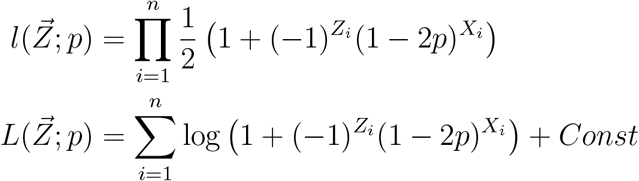

Taking the derivative to 0:

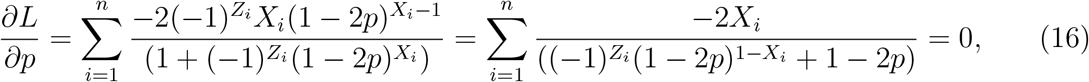

and division by 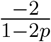 yields the solution.

Now, according to Le Cam’s theorem 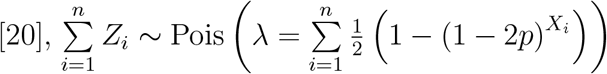 and the likelihood is therefore:

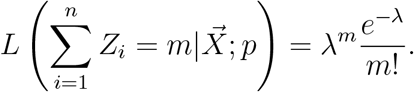

Now we look at the log-likelihood and take the derivative with respect to *p* to zero:

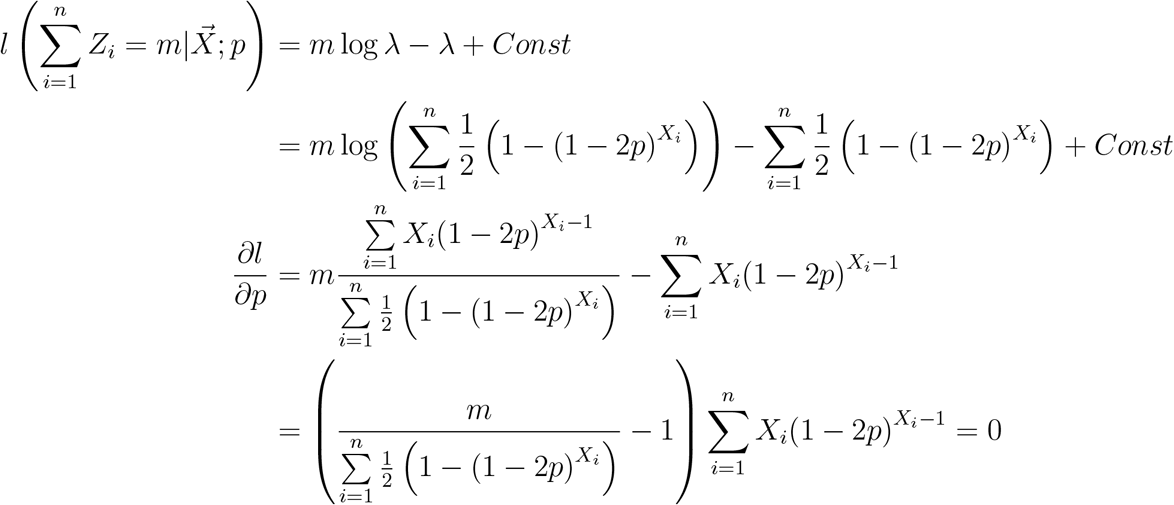

Leading to the solution:

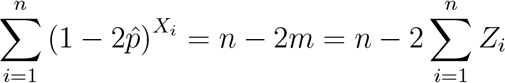

#### 1.5 Proof of Proposition 3

Let *λ*_*i*_ ∼ Γ(*α, β*), then the maximum a posteriori estimator of *p* holds:

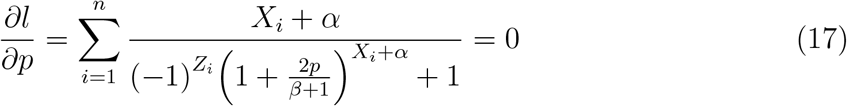

Subsequently, estimated values for *α, β* can be substituted for a numerical estimator for *p*.

*Proof*. We first compute the probability for each observation:

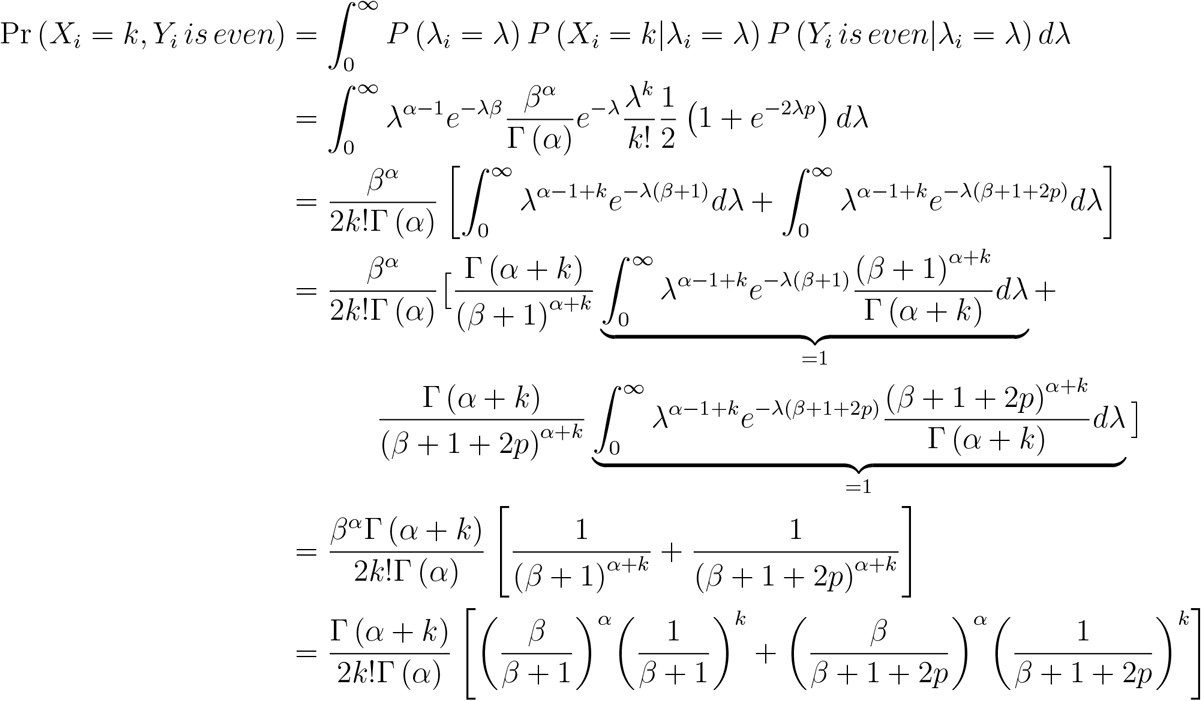

Hence, the likelihood is given by:

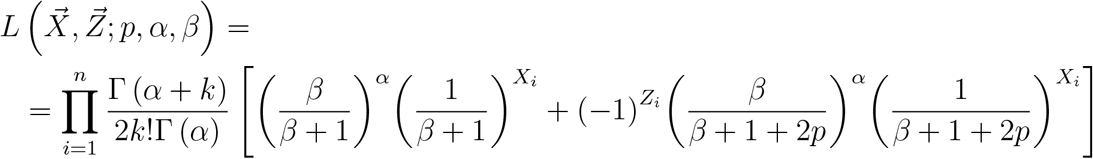

and the log-likelihood:

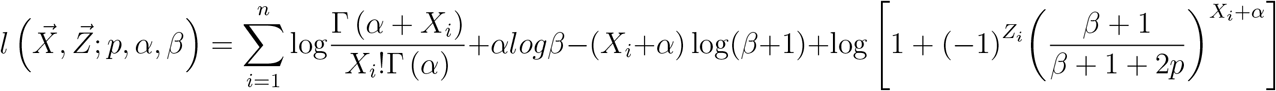

Now comparing the derivative with respect to *p* to zero:

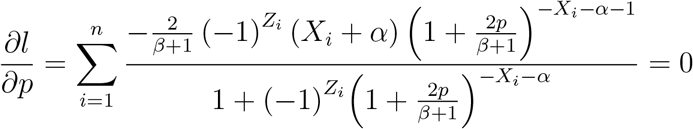

## 2 Simulation details

### 2.1 Phylogenetic tree simulations

The rate parameter for sites with no transitions along the tree is denoted as *E*, and we estimate it using the following simulation-based method. To generate *λ*, we use the following equation:

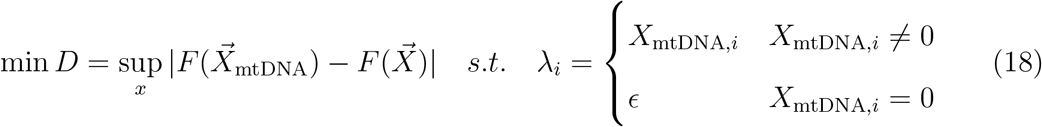

The value of *E* is chosen to minimize the Kolmogorov–Smirnov statistic. Figure 3 shows a simulation of *D*(*E*), with the mean of 1,000 runs for each *E* value. The minimum value of *D* is obtained for *E* = 0.0913 (marked in red).

**Figure 3:**
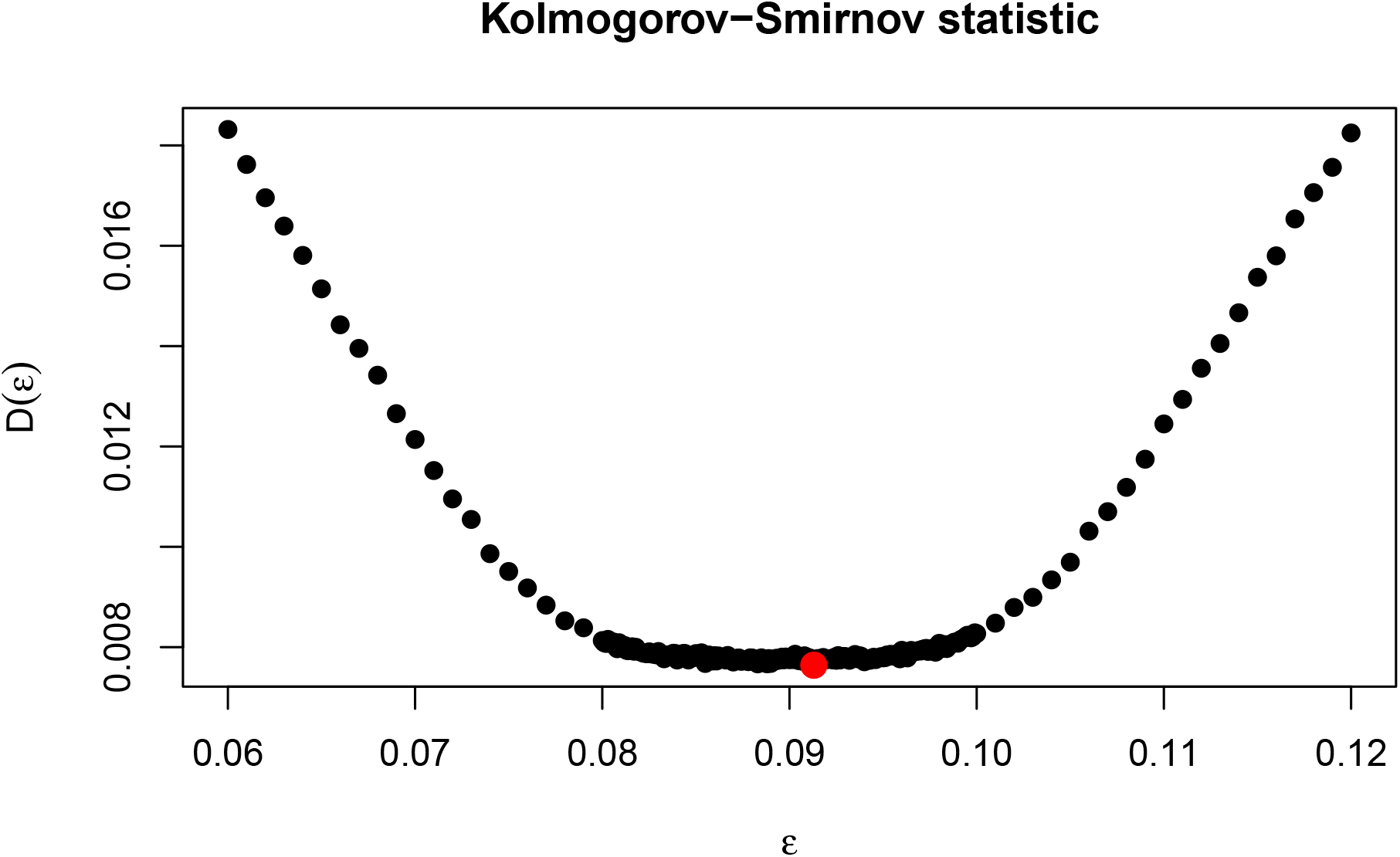
Kolmogorov–Smirnov statistic as a function of *E*. We performed 1,000 runs for each value of epsilon. The minimal *D*(*ϵ*) is marked red and equals *ϵ* = 0.0913.

To make the simulated data closer to the real data, we also model transversions. We estimate the transversion rate per site in the same manner as the transition rate, using the Kolmogorov–Smirnov statistic to account for sites with no transversions. This results in *ϵ*_transversion_ = 0.0149. To determine the nucleotide at a given site, we sample whether an odd number of transversions have occurred. If so, a random nucleotide is sampled from the two available transversion options. The resulting sequence is then input into BEAST2, but our methods still use only the sites without observed transversions. Finally, the analysis is limited to the gene regions in the genome (11,341 sites).

### 2.2 BEAST2 run parameters

The sequences used in this work were aligned using mafft [15], and the 11.3 kb of proteincoding genes were extracted and used for the analysis. The analysis followed the approach described in [34], where the best fitting clock and tree model for the tree were identified using path sampling with the model selection package in BEAST2 [14, 3, 21]. Each model test was run with 40 path steps, a chain length of 25 million iterations, an alpha parameter of 0.3, a pre-burn-in of 75,000 iterations, and an 80% burn-in of the entire chain. The mutation rate was set to 1.57 × 10E-8 and a normal distribution (mean: mutation rate, sigma: 1.E-10) was used for a strict clock model [10]. The TN93 substitution model [31] was used for all models. The tree was calibrated with carbon dating data from ancient humans and Neanderthals, where available [24, 10, 35], and modern samples were set to a date of 0.

## Footnotes

1 More precisely: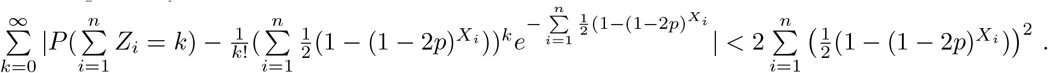

